# Compression through internal and external information bottlenecks resolves information intractability across brains, bodies and worlds

**DOI:** 10.64898/2026.09.24.751665

**Authors:** Arturo José Valiño, Axel Constant, A. César González-García, Laura Desirèe Di Paolo, Jadranka Verdonkschot, Felipe Criado-Boado, Andy Clark, Luis M. Martinez

## Abstract

Agents are constantly challenged to navigate uncertainty in a world that contains far more information that they can process, improving predictability by exploiting regularities from the environment. This frames the challenge of reducing information intractability as the main problem any artificial or biological agent needs to solve. We argue that this challenge is solved through the systematic use of information bottlenecks – a diverse suite of strategies functioning to compress and selectively filter environmental information so as to build tractable representations of the world.

We show that such bottlenecks are adaptive and operate across multiple scales, from the evolution of neural circuits and sensory systems to the actions that organisms use in real-time to select and structure their own sensory inputs. A key aspect of our proposal is that information bottlenecks are not only internal and biological, but can also be externalized through actions in the world. By transforming their environments, agents construct *external bottlenecks*—material and cultural mechanisms that reduce the cost of inference and transmit regularities that are shared across individuals and even across generations. From this perspective, niche construction, material culture and even social and cultural norms, can be also understood as adaptive informational mechanisms, collective information bottlenecks that alter our worlds in ways that promote predictive success.

Agents increasingly solve the intractability problem not only through evolutionary changes in their biology and internal neural mechanisms, but by systematically constructing and exploiting information bottlenecks in the world. Culture itself, we suggest, is just such a bottleneck - constructing mechanism.

**Significance Statement:** Understanding a system requires more than describing its structure; it also requires understanding the functions it serves and the operations it performs. In the case of human cognition, we propose the problem of Information Intractability as the operative guiding principle. That is, the world always offers far more information and possible actions than our limited brains, bodies and senses could ever process at once. From this guiding principle, we derive information bottlenecks as the solution to solve this challenge, by generating tractable compressed representations of the world that shape the coevolution of organisms and their environments.

This perspective offers a new lens for understanding biological evolution, niche construction, and cultural change as mechanisms that – at multiple scales of space and time – help solve the challenge of information intractability. Seen from this perspective, the human world emerges as a complex dynamic system in which agents continuously shape themselves and the environment to enable more precise and tractable action-guiding representations.

## Introduction

Any biological or artificial agent faces a fundamental computational challenge: the world contains far more information than any agent can ever fully process during behaviorally relevant timescales (1–5). This is the *challenge of information intractability* (CII): how can an agent discover useful regularities from an environment whose structure vastly exceeds its computational capacity?

Many disciplines across natural and social sciences concerned with perception, inference, and communication under uncertainty have implicitly or explicitly treated the CII. In neuroscience and psychology, concerns about the CII have motivated work on attention and working memory capacity (2, 3), and work on the extraction of structure from high dimensional sensory inputs (6–10). The Bayesian brain hypothesis (11, 12) addresses the CII by describing agents as approximate Bayesian inference machines that predict the states of the world with limited sensory evidence. Exact Bayesian inference is computationally intractable in general (13), and variational alternatives based on the computation of an approximate posterior have been proposed in response to these limits -- although approximation alone does not guarantee computational tractability (14, 15). In information and communication theory, the CII manifests as the problem of isolating informative signals from noisy channels (16–18), and in cybernetics, the CII underwrites works on the mismatch between environmental complexity and the limited response variety of any controller (19). Linguistics’ interest in the problem of the poverty of the stimulus and the need for inductive constraints in language acquisition (20, 21) is an expression of the CII, as are sociology and anthropology’s concerns with social institutions and cultural practices as collective strategies for reducing uncertainty and coordinating behavior (22–25). Views reflecting the CII in archaeology and cognitive anthropology include those arguing that material culture — artifacts, built environments, and symbolic technologies — function as transgenerational information storage media supporting cognitive processing by reducing environmental complexity (26–31). Further afield, certain art studies show that, through repetition and pattern, artistic styles encode a diversity of perspectives for representing the world (32, 33). Literary traditions likewise are thought to impose narrative forms that render complex social realities more tractable (34, 35), including the rules that guide oral transmission and literature (36). But to this day, to the best of our knowledge, no one has formalized an over-arching solution to the CII – one that would reveal the common features and functions uniting the many frameworks and disciplinary axes just described.

In this paper, we attempt to fill this gap. We introduce the framework of Tractable Compression (TC) through information bottlenecks as a means to formally express how information bottlenecks become the solution to solve the CII. The principle that guides TC is the reduction of the world into a tractable action-observation space (AOS) for inference. This action-observation space can be defined as an Information bottleneck.

Understood broadly, information bottlenecks compress and select smaller sections of information in the environment for posterior decoding in the brain (37). Every agent operates under such constraints and is, therefore, capable of sampling only a fraction of the available information from its environment. For example, human vision has been estimated to recognize only about 40 bits per second from roughly 20 MB of visual input entering the eyes each second, equivalent to extracting approximately two short sentences from an entire book of text (37).

We define Adaptive Information Bottlenecks (AIBs) as inherited or actively constructed information processing structures that compress patterns from the world into a tractable AOS and thus determine the signals a receiver can use to synchronize with, or understand the dynamics of the world. The brain, the body (e.g., sensory organs), and the environment (including artefacts) constitute an AIB, the structure of which allows a receiver to untangle regularity from an emitting process conveyed through a communication channel (Fig. 1A). Importantly, each of these components -- brain, body and environment -- can itself be understood as an AIB. Importantly then, bottlenecks are not biological constraints that limit information flow. Rather, they solve the intractability problem by selecting, compressing and limiting the information that an agent can use for inference.

**Figure 1.**
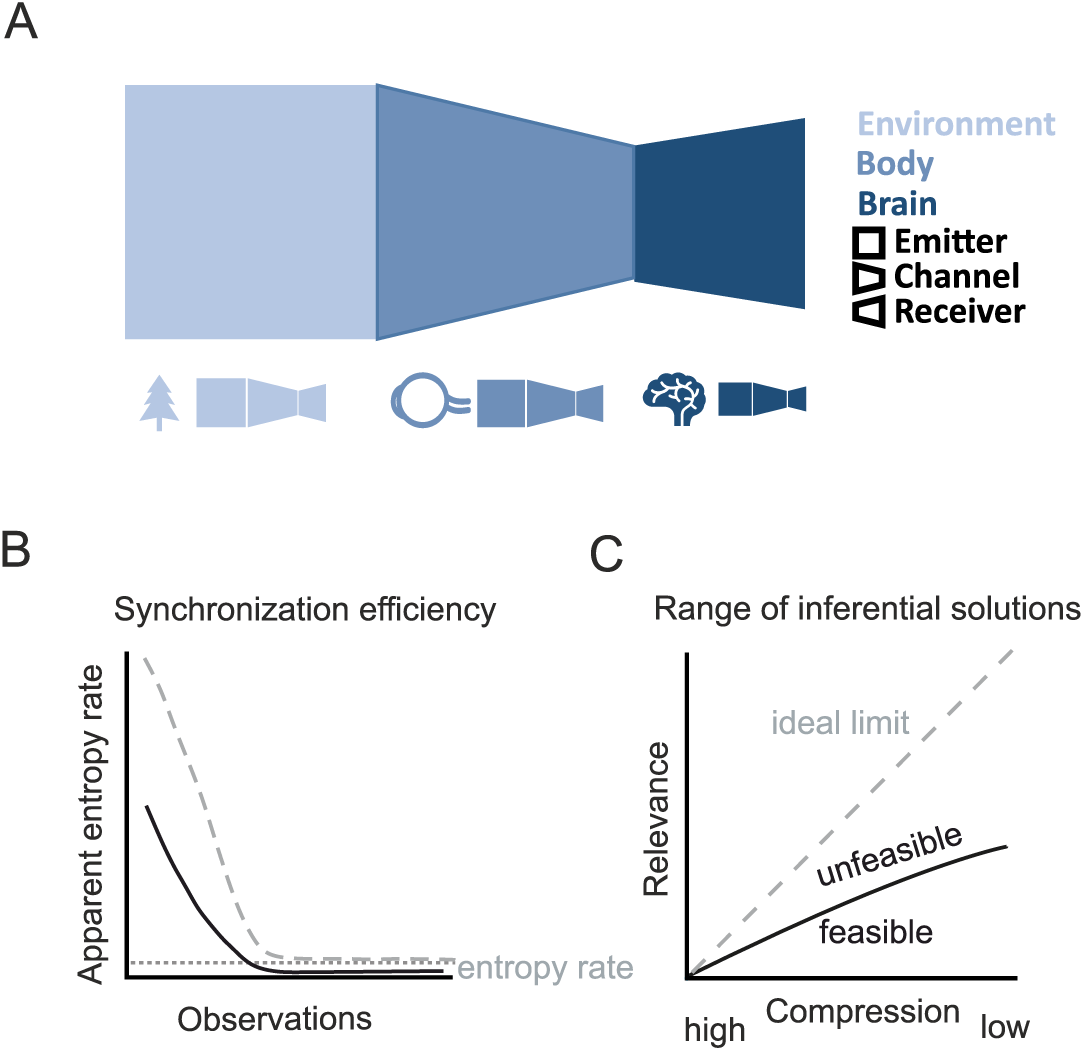
Adaptive Information Bottlenecks (AIBs). **A**. AIBs are made of an emitter, which emits a signal conveyed through a channel, and a receiver that can use that signal to infer its cause (the emitter). AIBs exist at all scales of organization within systems. **B**. Computational mechanics shows how AIBs constrain both the efficiency and precision with which an agent can synchronize to a process by characterizing how the apparent entropy rate converges toward the asymptotic entropy rate as longer observation histories are incorporated (dashed line). **C**. The information bottleneck method results reveal the range of inferential solutions that can be exploited under different AIBs. In both cases, results where compared reflecting the different ways in which an agent can approach the ideal limit of a fully resolving channel, in which there is complete access to all transitions relevant for prediction (dashed gray line in B and C).

We argue that AIBs are the force driving both the long-term evolution and short-term behavior of agents confronting CII. Further, the CII puts pressure on the class of inference procedures a receiver can implement to predict the emitter’s behavior, making each species or agent unique from an information theoretic perspective. The value of the TC framework, beyond its integrative potential, is thus to provide an explanation of the remarkable diversity of phenotypes observed in nature. These reflect the diversity of AIBs’ formed by cultural, evolutionary, and developmental processes.

Along the way, we provide a definition and typology of AIBs. Using simulation studies, we show the effect of different types of AIBs on the process of entanglement or synchronization (38) (i.e., on the amount of information required for solving the CII). We model AIBs as mappings from hidden-state transitions to observable symbols, and use the theory of computational mechanics (CM) (39, 40) to quantify how these mappings alter synchronization cost and precision. We then use the Information Bottleneck method (17) as a means to analyze the range of solutions that can be implemented by a population. The IB method (17) provides an abstraction of the compressed-relevance representations (41) available to non-ideal agents for a given AIB. Our framework, evaluates both results relative to an ideal AIB for a given emitter, providing a common reference point against which different AIBs can be compared. This allows to evaluate with the IB method (Fig 1. C), how close to the ideal agent different AIBs are, each representing populations with their phenotypic variations (i.e., space under the curve). We also explore the informational constraints imposed by an AIB and the tradeoffs in cost and precision respect to the ideal limit (Fig 1. B). We explore the fundamental informational tradeoff between the size of channel capacity, the different AOS mappings imposed by the AIB and the efficiency with which one can represent the cause of sensory signals, equivalent to the trade-off between reading the whole book to get the story straight and skimming the pages at the risk of missing out on important hidden meanings.

We conclude with a broad discussion of the TC framework, focusing on its explanatory role as a driver of both cultural and biological evolution. AIBs help explain: (1) the evolution of the architecture and connectivity of the nervous system; (2) how concepts, in particular, and language, in general, can be understood as forms of internal AIB that mitigate the cognitive effort of synchronizing with the environment; and (3) how agents actively solve the intractability problem by offloading computation to the environment, creating externalized bottlenecks that reveal new hidden states and select relevant observations for behavior. The upshot is a unified formal framework that shows how biological form, active niche construction, and the distinctive effects of human culture combine to allow us to make sense of the world.

## Results

### The Tractable Compression Framework

Any organism faces the challenge of solving the CII. To do so, agents must compress the world into a tractable AOS that enables to efficiently untangle randomness from the world. The result is a compressed reflection of the world’s hidden dynamics -an AIB-that restricts the information available for inference while enabling the exploitation of its hidden regularities.

AIBs thus determine the subset of the information space an agent has access to. We call these bottlenecks adaptive because they are shaped across timescales to make prediction tractable. Some are inherited and relatively fixed during online behavior, such as sensory architectures; others are actively modified through learning, attention, action, or environmental construction (e.g., human-built environments). Thus, we put forward a view of AIB as processes that are a reflection of their information space, that condition the information that can be accessed to decode regularities from the world and that constrain the inference strategies each species or agent can develop.

Let the information space be the set,

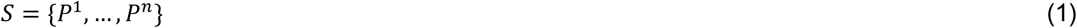

of all subprocesses that constitute the environment, defined as transition dynamics over states A, B, C, D…. An agent selects a sensing subset

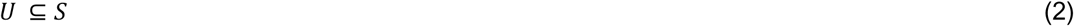

where the synchronization with U is achieved through a derived information space, S’, the AIB (Fig. 2 A, B). The purpose of S’ is to mirror the information of S (the full information set) and to compress the observations related to the transition dynamics of *P_j_*, so as to define the channel capacity (C) of the AIB.

**Figure 2.**
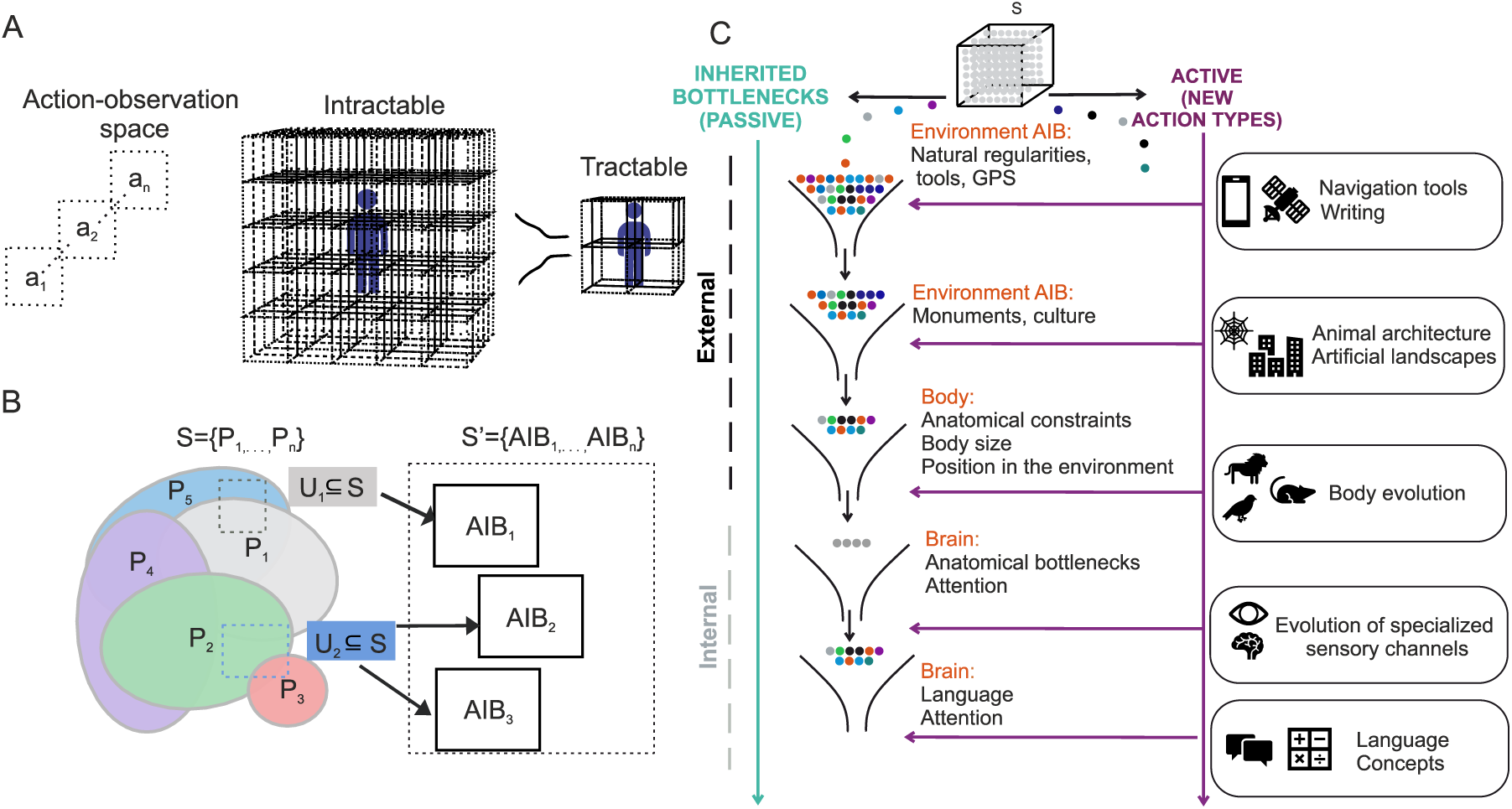
The tractable compression framework. **A.** An adaptive information bottleneck (AIB) transforms an intractable action–observation space into a tractable one by restricting the diversity of actions used for synchronization. **B**. The perceptual landscape S contains multiple subprocesses. The selected sensing subsets U1, U2 are accessed through a mirror process S’. Different AIBs structure access to different sensing subsets, and more than one AIB may also constrain the same subset. The AIB addresses the intractability of the full perceptual landscape by selecting a subset U⊆S and constraining the action repertoire through which observations can be revealed. **C**. AIBs may arise at world, body, and brain scales. They may be passively inherited, such as anatomical constraints or environmental regularities, or actively constructed, such as tools, maps, attention, concepts, and learned sensorimotor strategies. Color coded are diversity of action-observations from S’ to access de information from the total perceptual space S. At each AIB bottleneck step new actions-observation types can be used.

Our AIB builds from the measurement-channel perspective of computational mechanics derived from Shannon’s information channel (for a more in-depth review of these concepts see (39, 40). In this sense, the AIB is described as a process (emitter), a measurement/symbol stream or channel, and a receiver. However, we consider actions to determine the mapping of the state transitions that can become observable though the process. Therefore, we propose to work with an action-measure-channel scenario by assuming that *actions generate updates in the observation channel* (Fig. 3). Because observations are generated by the transition dynamics over states (e.g., emitting 0 as *P_j_* transitions from states A to B), a receiver can discover the transition dynamics by making observations. For the sake of simplicity, we assume that making an observation is equivalent to acting, where acting involves defining the symbols or signals a receiver will use to reveal the transition dynamics. This means that channel capacity is expressed as:

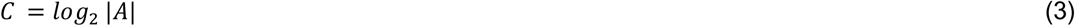

**Figure 3.**
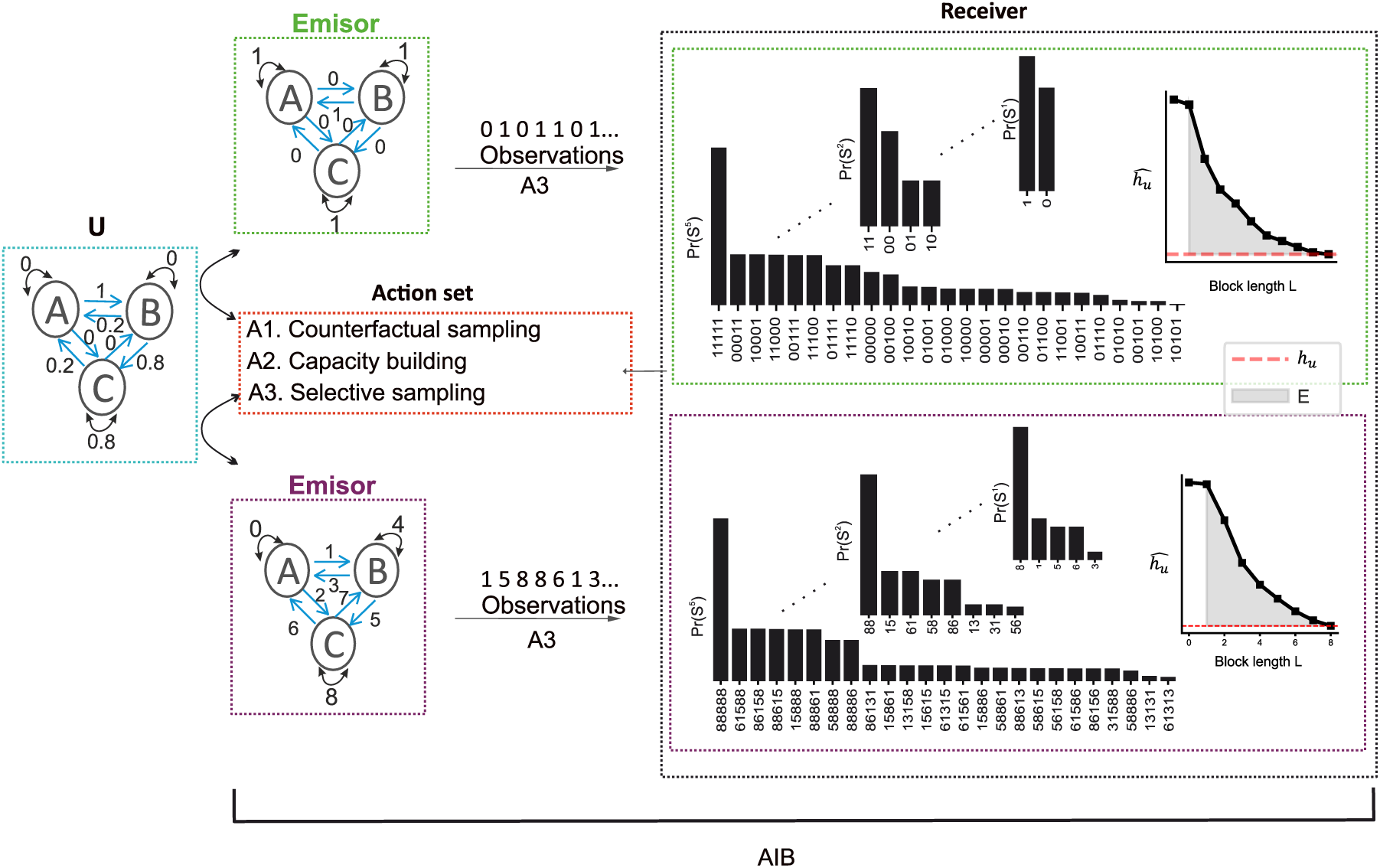
Tractable Compression model. An AIB entangles with a subset of the information space (U). There are 3 types of possible actions that an AIB can implement: **(A1)** Counterfactual sampling: where the symbols that reveal hidden state transitions are permuted. **(A2)** Capacity building: where new symbols that can be emitted by the process become available within the signal stream. **(A3)** Selective sampling: determines the transitions that are revealed to generate the measurement stream of observations. In the receiver, the effects of these actions were computed using block entropy. **Left**: Block entropy for different L values (blocks of observations as seen in the black histograms). **Right**: Entropy rate as function of L (excess entropy in shaded gray).

where A is the diversity of actions, which is equivalent to the diversity of observations, or signals an agent can use to understand a process. Thus, the discrete diversity of different actions the agent can use to receive observations determines the size of the alphabet (A) and, therefore, the channel capacity C, where an action is defined as the symbol an agent uses to map a hidden state transition.

For instance, out of all the hidden state transitions in *P_j_*, the eye as an S’ process will mirror and sample an approximation of these transitions with a partial representation determined by retinal image size (e.g., a small eye may resolve only 10 distinct symbols, whereas a larger eye may resolve 20), thereby limiting the channel capacity (e.g., limiting C to *log*_2_ |10|, which would have been otherwise *log*_2_ |20|).

One can imagine 3 types of possible actions that an AIB can implement:

(**1**) counterfactual sampling, where the symbols are permuted while preserving their original order (e.g., “what if” the signal 1 in the string 1 2 0 1 emitted by the A to B transition instead was a 2, such that the whole string becomes 2 2 0 1?). This preserves the same channel capacity C, but makes the system sensitive to different transitions.

(**2**) capacity building, where new symbols that can be emitted by the process become available within the symbol stream (e.g., 0,1,2,3 instead of 0,1, and 2);

(**3**) selective sampling, where, as in the example above, only some combinations of symbols within an observation stream are selected (e.g., 1, 2).

In contrast to the standard Information Bottleneck notion –-where the bottleneck is a compressed representation derived from any signal respect to another (17), or the selection of relevant information from the environment (37)--our formulation conceptualizes the bottleneck as a distributed system encompassing the process, emitter, channel, and receiver, in which compression, selection, and capacity are jointly shaped by actions across all components. This implies that actions are not properties of any single element of the AIB (i.e., the emitter, channel, or source), but rather a capacity of the whole AIB to transform the signal source. The receiver can then use any inference strategy of choice.

### Why do AIBs solve TC?

A completely random environment will be impossible to navigate. Because natural processes have regularities (42), hidden regularities in S, the mirror process is able to exploit them and untangle them from randomness. Those regularities that cannot be exploited remain as apparent randomness (39). Thus, the agent never interacts with the whole complexity of the word, it interacts always through an AIB that compresses the information space making synchronization tractable. Tractable compression therefore states that agents interact with a subset of the perceptual space U through a mirror process, that reflects and makes tractable the underlying information from the world.

As an example, let us consider the problem of visual navigation through a landscape. In this case, an agent has to solve the CII related to building precise egocentric and allocentric representations of the world (43). This is achieved through coupled AIBs across environmental, bodily, neural, and cultural systems (Fig. 4 and SI for detailed example), that enable the agent to progressively untangle more and more regularities from the world. Beginning with the agent’s position in the environment, which already constrains the action-observation space, successive bottlenecks emerge through technologies such as GPS, sensory organs, neural circuits, and cultural artifacts. These AIBs operate across multiple temporal scales to selectively compress information while preserving task-relevant structure, progressively transforming an intractable inference problem into a manageable one.

**Figure 4.**
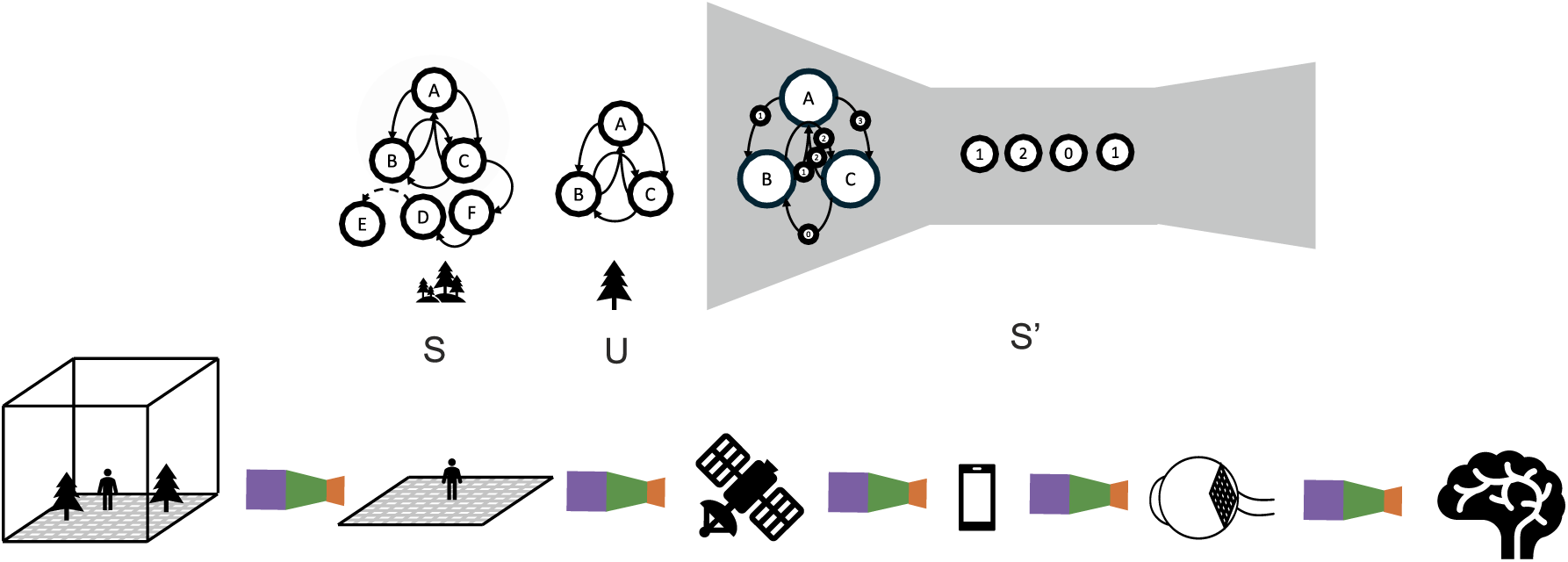
Coupled information bottlenecks enable the construction of tractable spatial representations during navigation. Top: schematic illustration of an AIB that mirrors the informational structure of the environment by transforming an initially intractable action-observation space into a tractable representation. Bottom: successive bottlenecks involved in visual navigation. The agent’s position in the environment constrains the available action-observation information; satellite signals further reduce uncertainty about location; ocular optics limit the sampling of visual information reaching the retina; retinal circuitry compresses photoreceptor signals into a reduced neural code transmitted through the optic nerve; and subsequent neural and cognitive processes generate behaviorally relevant representations for navigation. Cultural artifacts such as maps, roads, landmarks, and navigation devices can function as higher-order bottlenecks that further simplify inference and action. Together, these coupled bottlenecks progressively make spatial navigation computationally tractable.

In efficient coding (44), compression removes redundancies through the exploitation of statistical regularities of the world to maximize the mutual information from stimuli and neural response due the limited channel capacity of the nervous system (45–47). Our notion of compression is different. It is not simply the elimination of redundancies, but the construction of a tractable process capable of detecting and exploiting the regularities that characterize the information space. Compression, in this sense, refers to the formation of an AIB that extracts the amount of redundant and non-redundant information required for precise and tractable inference. Thus, compression should not be understood merely as redundancy reduction. Rather, it is the emergence of a process that preserves the structure necessary for tractable inference. An AIB thus construed compresses an information space by: (1) selecting a sensing subset and, (2) by determining the states and actions used to reveal the hidden transition mappings.

### Passive and active AIBs

We can begin the task of classifying different types of AIBs by considering the passive or active nature of the action operated by the channel. Selective and counterfactual sampling are “passive” actions as they keep the channel capacity fixed. This may be the case, for instance, of inherited, and developmentally stable sensory organs that compress observations with a fixed channel capacity on the timescale of the current task. As an example, in the retina, the joint actions of horizontal, bipolar, amacrine, and ganglion cells actively compress the observations that reach photoreceptors, within a relatively fixed anatomical and physiological channel capacity (48, 49). By way of contrast, active bottlenecks are constructed when agents introduce new action–observation types that increase the channel capacity of the AIB. For example, a GPS system may increase its number of satellite signals to better infer position.

Note that AIBs are integrated processes that constantly switch from a passive and active cycle due to actions that happen at different temporal timescales. A bottleneck may be actively shaped over long evolutionary or developmental timescales, yet function as passive during the short behavioral timescales of perception and action. The retina illustrates this point. Although retinal circuitry is actively shaped over longer biological timescales, it typically functions as a passive bottleneck during moment-to-moment vision.

### Internal and external AIBs

A core aspect of our framework is that we further distinguish AIBs according to the domain in which the channel is realized, that is in the internal or external domain (Fig. 2C). Note that here, the internal/external distinction is meant to capture the emitter/receiver relation. But AIBs occur at all scales of organization -- including within the brain. Internal AIBs may connect a sensory organ (i.e., an emitter), an early region of the brain (i.e., channel), and a higher region of the brain (i.e., receiver). This delivers a partial reconfiguration of the nervous system that can expand channel capacity, often at the expense of reducing capacity elsewhere in the nervous system (50–52). At a more abstract level, we consider concepts and language as actively built information bottlenecks created at one temporal timescale that offload the cognitive effort required to passively understand the complex dynamics of an information space at another temporal timescale (Fig. 2C). Attention can likewise be understood as a set of temporally shifting active and passive bottlenecks that dynamically filter the action-observation space (AOS). However, whereas language compresses and structures information into reusable conceptual forms, attention dynamically regulates access to the perceptual information in the first place (53).

External AIBs, for instance, include the physical body -- from the point of view of the brain -- where the body is understood as a channel filtering and simplifying the sensory information (i.e., observation) to be processed by the brain. This view, common in embodied cognitive science is nicely illustrated by passive dynamics walking systems (54, 55), which greatly simplify the computations needed to generate bipedal walking though limiting the degrees of freedom of movements and letting gravity along with physiological constraints control the walking motion.

External AIBs capable of playing this kind of role include aspects of material culture, (e.g. way-signs), technologies (e.g. GPS), and social institutions (e.g. stock markets) all of which constrain and create action–observation spaces. External AIBs are modifications of the world that stabilize useful action–observation mappings, making future synchronization more tractable by reducing the computational cost of selecting and compressing relevant observations. In this sense, the evolution of cortical areas reflects not only the acquisition of new perceptual capacities, but also the reduction in flexibility that follows when the channel capacity of some bottlenecks is expanded at the expense of others (56). As mentioned, socio-cultural analogues occur in tools, notations, and institutions, which similarly enable certain actions by constraining the range of viable alternatives (57).

Across both cortical organization and human culture, tractability is achieved through the stabilization of neuro-ecological artifacts that make some forms of action and prediction easier while making others less accessible.

### The Tractable Compression Model

We built a model to explore how the AIB determines the ability of an agent to synchronize with a hidden process. We adopt an action oriented computational-mechanics perspective focusing not on the agent’s inference strategies but on the structural limits imposed by the bottleneck itself. This allows us to explore the limits on synchronization efficiency and precision given the diversity of actions imposed by the AIB (Fig. 3). In addition, we use the IB method (17) to explore the range of possible inferential solutions an agent could adopt given the inherited AIB.

To isolate these limits, we fix an emitter process of 3 interconnected states (A, B, C) that is, in principle, capable of mirroring the hidden dynamics of the underlying process when sufficiently informative observations are available (Fig. 3 and 5). We then vary only counterfactual and capacity building actions. This allows to characterize how the action repertoire shapes the observation channel and constrains synchronization efficiency and precision. We deliberately abstract away selective-sampling strategies, since our aim is not to identify the optimal inferential architecture, but to show how the AIB sets the precision and efficiency of synchronization.

**Figure 5.**
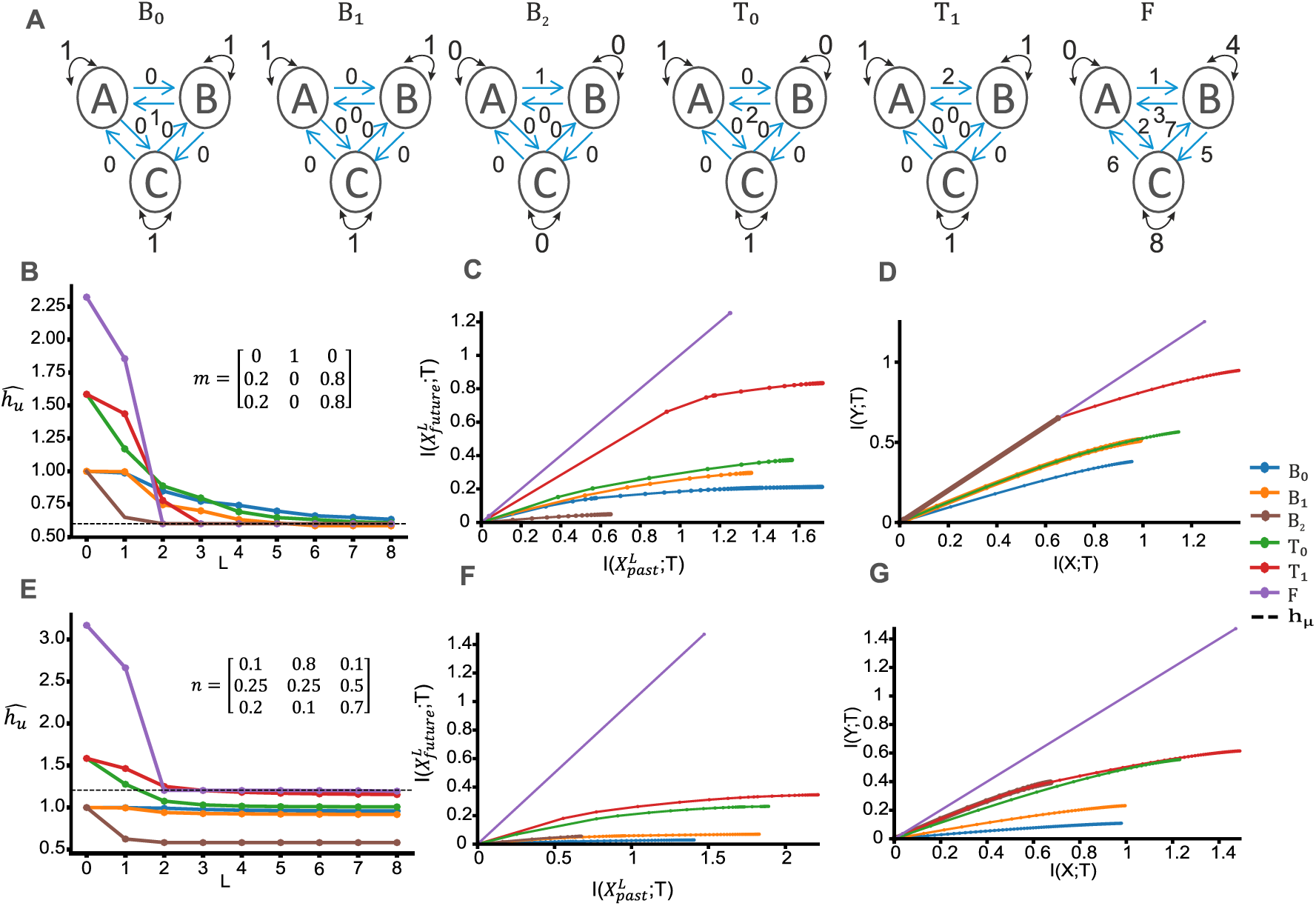
AIBs constrain the efficiency and precision of synchronization with the world. **A**. Different simulated AIB mappings due to changes in channel capacity or counterfactual sampling condition synchronization efficiency and estimation. **B**, **E**. Estimated entropy rate 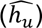 for the m and n processes. Dashed line in B and E corresponds to the true entropy rate h_μ_ of the process (see methods). **C**, **F**. Information Bottleneck trade-off between compression and predictive relevance for future 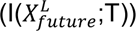 and past information 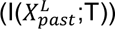 across AIB channels, shown for the simple (m) and complex process(n), respectively. **D**, **G**. Information Bottleneck trade-off the observation channel *I*(*X*; *T*) and hidden state process *I*(*Y*; *T*) across AIB channels. Across processes, increasing channel capacity and counterfactual sampling expanded the relevance-compression profile, but channels differed in how efficiently this additional capacity was converted into predictive relevance. The all-transitions channel represents the limiting case, because the target process is deterministically specified by the observation stream.

### Computational mechanics

Computational mechanics (CM) (40) provides an information-theoretic framework for quantifying the dynamics of uncertainty itself, revealing how randomness and structure unfold as an “ideal observer” accumulates observations. The ideal observer is a computationally unconstrained agent with direct access to all possible noiseless observations, whose task is to disentangle structure from randomness (39).To synchronize with a process is, therefore, to describe how random the source is. A process has both apparent and real randomness, and regularities and structure that cannot be untangled remain as apparent randomness (39).

In CM, the randomness of the process is quantified through the growth of a special quantity, *block entropy*. That is, the change in entropy as more blocks of observations are added. This, in turn, reveals the hidden regularities and quantifies the information required for prediction:

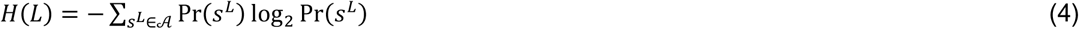

where Pr(*S^L^*) stands for the probability distribution over blocks of L consecutive observations.

Thus, *S^L^* = *S*_0_*S*_1_, …, *S*_*L*−1_ stands for a particular block symbol of consecutive environm ent observations of length L, where each symbol *S_i_* ∈ *A*. Here *A* is the observation alphabet, that is, the set of all distinct observation symbols that may be emitted. The sum goes over all possible blocks of L consecutive symbols. The scaling of H(L) with L (Fig. S1) underlies several information-theoretic quantities that characterize both the randomness and the statistical structure of the process (Fig. 3 and 5).

These quantities are, the entropy-rate (h_μ_), excess entropy (E) and transient information (T). In this framework, h_μ_ captures the residual randomness of the observed process, E the cost of amnesia, that is, the amount of past information I need to keep track of h_μ_, and T quantifies the cumulative information cost required for synchronization. Thus, these measures allow us to compare how different AIBs alter both the precision and efficiency with which an observer can synchronize with the same hidden process. Full definitions and estimation procedures are provided in methods.

### The simulation

We simulated two different three-state hidden Markov processes to compare the true entropy rate of the process (h_μ_) with the apparent entropy rate (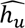) imposed by the AIB. This corresponds to the ideal synchronization an agent can achieve constrained by the AIB (see methods to see how quantities were computed). The simple process (s) allowed all channels to approximate the true entropy rate. The complex process (c) required an increase in channel capacity to avoid underestimating the true randomness of the process (Fig. 5).

We compared six observation channels differing in both their alphabet capacity and the way hidden transitions were mapped onto observable symbols (Fig. 5A). Three Binary channels (B₀, B₁ and B_2_) differed in the mapping between transitions and symbols—counterfactual sampling. Two Ternary channels (T₀ and T₁) increased alphabet capacity by one symbol but differed in whether the added symbol captured an informative or uninformative transition –capacity building and counterfactual sampling. The Full transition (F) assigned a unique symbol to each distinct hidden-state transition. Thus, differences in the block entropy profiles came entirely from the different transition to symbol mappings caused by the counterfactual sampling and capacity building actions. Together these results show how different AIBs mappings shape synchronization efficiency and precision.

### AIBs determine inference efficiency and precision of synchronization

In this simulation of the simple process, the hidden process remained unchanged, with a true entropy rate ℎ*_μ_* = 0.601 (Table S1; Fig. 5B). The initial binary channel (B₀) apparent entropy-rate 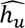, overestimated h_μ_ and produced a relatively high excess entropy E and transient information T of all the binary channels, indicating that a larger amount of information and observations history was required to untangle uncertainty and synchronization.

Refining the binary mapping without increasing capacity improved performance. In B₁, reorganizing the same two-symbol alphabet reduced E, and T, while improving the entropy-rate estimate 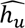. The most efficient mapping was B₂. It assigns a dedicated symbol to a single highly informative transition (A to B) and groups all other transitions under the second symbol. This produced a very low T and rapid convergence, outperforming all other channels despite its minimal capacity.

The ternary conditions—T₀ and T₁-- reveal that increasing capacity is only beneficial when the added symbol is informative. The uninformative expansion T₀ improved the entropy-rate estimate 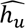, but retained high E and T. In contrast, placing the added symbol on an informative transition produced a very good approximation to ℎ*_μ_* and a low T value. The F channel yielded the most accurate 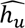, but also produced the highest E and larger T than T₁. So, a richer observation alphabet untangles more of the process structure but also requires more information for synchronization.

Overall, these results show that synchronization efficiency depends not only on channel capacity, but on how transitions are mapped to symbols.

### Effective changes in AIBs’ channel capacity are necessary for accurate synchronization

We next examined a second process (c) with more complex dynamics where the synchronization efficiency changed across channels (Table S2; Fig. 5E). In this case, the binary channels that performed well in the simple process now underestimated ℎ*_μ_*. The ℎ*_μ_* of the process exceeded the capacity (C) of a binary channel. Thus, refining the mapping within a fixed binary alphabet was no longer sufficient. As a result, capacity increases are necessary. However, efficiency depends on how the added symbol was assigned. The uninformative expansion (T₀) provided little improvement and became inefficient once information processing costs are considered. By contrast, the informative expansion (T₁), produced a more accurate 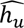 and a more efficient synchronization profile. The fully resolved channel (F) yielded the most accurate estimate of the entropy rate, but had a larger synchronization cost -- a higher T value. Together, these results show that accurate synchronization requires an increase in channel capacity when the entropy rate of the process exceeds the representational limit of the original alphabet. However, this increase is beneficial only when the added symbol resolves an informative transition. This identifies an efficient intermediate regime in which a targeted increase in C improves synchronization without incurring the additional cost of a fully resolved channel.

### AIBs determine the range of possible solutions imposed by the compression-relevance tradeoff

We have shown through computational mechanics how the AIB determines the ability of an agent to synchronize with a hidden process. This defines the ideal limit of the predictive structure imposed by the actively built or inherited AIB—a receiver that can use all sampling actions with no constraint. Hence, the entropy rate of the observable process defines the amount of randomness that is accessible to an ideal observer under a given AIB.

We now extend this approximation using the IB method (17, 41, 58, 59) to explore the range of compressed-relevance solutions that can be exploited by non-idealized agents within the constraints imposed by the AIB. This relevance compression tradeoff quantifies how much predictive relevance can be retained as observations are progressively compressed into lower-dimensional representations (Fig 1, see SI for further details). Thus, the IB method is not used here as a literal selective sampling mechanism, but as a characterization of the different inference procedures that could be exploited within the subset of action types made available by the AIB.

We, therefore, estimated the information bottleneck (IB) relevance-compression trade-off (41, 58, 59) for the different AIBs. The trade-off was computed in two different ways: (i) compressing the observation stream to preserve predictive information about the future 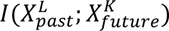 and (ii) compressing observations with respect to the hidden states of the generative process (see methods and SI). In this sense, the IB does not define the AIB. It explores what a non-ideal predictive agent could do with the constraints imposed by the AIB. Across channels, this analysis confirmed that the range of useful relevance-compression solutions is determined by the AIB. Increasing channel capacity expanded the relevance-compression profile, but the magnitude of this improvement depended on whether the additional symbols preserved distinctions relevant to the target.

### AIB trade-off between the past and future information of the channel

The results show that for the simulated simple and complex processes, an increase in channel capacity is always beneficial, although channels can have more efficient capacity increases, enabling a better relevance-compression profile (Fig. 5 C, F). In addition, the refinement of observations leads to an improvement in the compression-relevance trade-off. Furthermore, the channels exhibited different efficiencies relative to the information they contained when observing the normalized profiles (Fig S2. A, C), excluding the Full transition (F) channel. For the simple process, the channels behaved similarly, and for the complex process, there was more variability, with the coarser channel being extremely efficient with the information it had. Finally, the F channel is uniquely efficient because the future (Y), at a macroscopic scale, is a deterministic function of the observation stream (X) (Y=f(X)), and thus the compressed representation Z is equally informative for X and Y (I(X,Z)=I(Y;Z), see (60)).

### AIB trade-off between the observation channel and hidden state process

Our results show that increasing the channel capacity does not uniformly improve performance. Instead, we observed that some increases in capacity were inefficient for both the simple and complex processes (Fig. 5 D, G). This reflects the exploration cost of sampling uninformative states in the AIB framework. Furthermore, improvements in the observation stream without changes in capacity can lead to better profiles. In addition, the behavior of the F channels remains highly efficient, as the observation channel is a near-deterministic representation of the hidden process (60). However, the behavior of the coarse channel shows clear differences: in the simple process, it becomes highly efficient relative to the information it contains, aligning with the performance of the fully resolved channel until it reaches its limit. In contrast, in the complex case, it aligns with the profile of the efficient capacity increase channel until it reaches its limit.

In the efficiency plane, for both processes, a properly oriented increase in channel capacity is very efficient. The F channels represent the ideal scenario where there is no trade-off, and the rest of the channels have a similar efficiency (Fig. S2 B, D). However, for the simple process, the coarse channel exhibits a slope close to one. Thus, in this case, the AIB has already imposed an optimal compression with respect to the hidden process such that no further information can be removed without compromising its relevance (41).

## Discussion

Tractable Compression (TC) proposes that the fundamental computational problem facing any biological or artificial agent is information intractability, and that adaptive information bottlenecks (AIB) are the solution, making the world computationally tractable. Therefore, agents never interact with the full complexity of the world. They synchronize through an AIB, a compressed reflection of the world that constrains their actions, and therefore, the information that is accessible to solve the challenge of information intractability (CII), making each agent, species or group—depending on the scale of analysis-- a unique information processing machine. Thus, we define an AIB not as a biological constraint that limits information flow, but as the solution to the CII. From this perspective, AIBs drive action across evolutionary and developmental timescales, while phenotypic diversity reflects the variety of solutions that agents can adopt to solve the CII. Variability in morphology, cognition, and behavior can therefore be understood as alternative strategies for making different aspects of the world tractable.

We propose that there is a selective pressure in evolution to build AIBs that expand effective channel capacity (understood as the diversity of actions available to sample the environment) as well as reducing the amount of information required for synchronization to solve the CII. Our TC framework proposes how these processes are inherited and actively built across timescales to determine how efficiently and precisely an agent can synchronize with the external world with respect to an ideal limit. Following our results, computational mechanics shows how AIBs constrain both the efficiency and precision with which an agent can synchronize with a process. In turn, the Information Bottleneck analysis reveal the range of inferential solutions that can be exploited under these constraints. Together, these results show that changes in AOS mappings due to capacity building and counterfactual sampling generate different synchronization schemes, determining both how effectively an agent can synchronize with a process and how much predictive relevance can be extracted from it. In both cases, we compare results respect to a fully resolving channel (F), in which there is complete access to all transitions relevant for prediction, the ideal limit. This is represented by the deterministic boundary observed in the IB trade-off, and in the transient information (T) in CM. From this perspective, the CII generates selective pressures favoring AIBs that more closely approach this ideal limit.

As with many evolutionary problems in the life sciences, questions surrounding the CII can be organized into two complementary explanatory levels: (1) proximate explanations, concerning the mechanisms by which a trait operates, and (2) ultimate explanations, concerning the evolutionary processes that account for why the trait exists (61). Under the CII, a proximate question would be: How do agents extract actionable regularities from environments whose combinatorial complexity vastly exceeds its computational capacity. In turn, an ultimate question would be: Why did the mechanisms that support information tractability evolve the way they did, given the fundamental challenge of computational intractability. From this perspective, computational intractability exerts an inescapable selective pressure on the evolution of cognitive systems. Evolution should therefore be expected to favor anatomical, physiological, behavioral, social, material, and cultural traits that reduce the operational complexity of inference by exploiting, internalizing, or constructing stable regularities (i.e., AIB) in the environment. Evolution can thus respond to the CII in three different ways. First, by improving the computational apparatus by generating better brains, sensory systems, memory capacity, attentional mechanisms, etc. Second, by exploiting the statistical structure of the environment by selecting better priors, heuristics, etc. And, third, by modifying the environment itself so that inference becomes easier, resulting in a world that is more computationally tractable for the agents inhabiting and transforming it. There is a driving force guiding every agent towards untangling organism-relevant randomness from its environment, and thus we should be able to frame the evolutionary history of many traits as being geared towards solving the CII.

Our results suggest that AIBs should be constructed so as to approach the ideal limit through “action” (i.e., structuring of channel capacity) at different timescales. At evolutionary timescales, we can see how body and brain architectures adapt to (and make available) different information spaces. At developmental timescales, we see how actions shape the channel capacity and observations used for prediction. But actions that increase channel capacity and compression have a cost, so there is always a tradeoff: selecting a region of the signal normally comes at the expense of decreasing capacity on other regions (50–52). We conclude by discussing some implications of our framework for biological and cultural domains, showing how the TC framework can be used to (i) integrate disciplines across the natural and social sciences, and (ii) provide an explanation -- in proximate and ultimate terms -- of the evolution of traits in response to the CII.

### Internal Information bottlenecks

#### The visual system

The visual system is an accurate example of an AIB in biological evolution. Eye size increases with body size and larger eyes promote better visual acuity (62, 63). The increase in retinal image size–channel capacity– due to larger eyes - implies an increase in visual acuity, understood as the ability of an animal to resolve static spatial details (64, 65). Furthermore, visual acuity also scales with primary visual cortex size (V1) further enhancing acuity as processing capacity in higher stages is increased (66, 67). More generally, increases in neuron number are expected to expand the computational capacity of neural systems (68, 69).

Nevertheless, there is always a tradeoff – a metabolic cost – when increasing capacity in an AIB. Acuity is increased at the expense of reducing the diversity of sensors that tile the retina (70). Furthermore, the topological disposition of these sensors varies across species. Given this diversity of sensors, evolution has reached different counterfactual sampling solutions by changing how these sensors are tiled in the retina across species. Simple examples include the visual streak that oversamples the horizon in wolves, the fovea of carnivores and primates (71), and the overrepresentation of specific types of quick responding neurons in the ventral part of the mouse retina that point to the sky (supporting quick detection of looming behavior from an aerial predator) (72).

This tradeoff in capacity is also evident in changes to the relative size of different brain areas: changes that reflect the varying ecological needs and lifestyles of different species. There, channel capacity is preferentially increased through the differential allocation of neuronal and metabolic resources across cortical systems, constraining the resources available to alternative processing pathways (51, 52, 73, 74). Carnivores and primates, for example, have larger visual areas compared to their somatosensory regions, whereas species like mice allocate their neural resources to larger somatosensory areas, reflecting their reliance on tactile input (56). This tradeoff in channel capacity is even evident on short timescales AIBs that respond to rearing environmental conditions by altering sensory cortical size relationships on developmental timescales. In this sense, the evolution and development of cortical areas reflects not only the acquisition of new perceptual capabilities but also the increasing commitment of neural resources to particular information channels, which may reduce the flexibility available for alternative computations (52).

Tradeoffs in channel capacity also manifest at other macro scales of neuronal function. Animals with a higher degree of binocular view have higher visual acuities at the expense of reducing the overall size of their lateral vision (75). Thus, changes in channel capacity and counterfactual sampling determine acuity which limits the distance at which observations can be properly resolved, shaping the inference strategies of species that live in a small haptic world by favoring quick detection mechanisms (mice) or in a big visual world (carnivores) by favoring strong visual correlations and precision In conclusion, neuro-ecological artefacts such as the retina and the visual cortex are clear examples of how such artefacts coevolve with the environment. This reflects the diversity of preferred actions given differing entanglements with an ecological niche. In humans, Material Engagement Theory (76) suggests that similar principles of co-adaptation continue at the cortical level, where long-term engagement with tools turns the brain itself into a genuinely bio-cultural artefact.

### Externalized Information Bottlenecks (EIBs)

Agents do not only decode regularities from the environment–they also create them. Through action, agents reshape the world, producing externalized information bottlenecks (EIBs) that reveal new hidden state transitions and compress the most relevant observations of a process. These changes in the environment, if done optimally, should generate AIBs that approximate the true entropy rate of the process and decrease the information required for synchronization – the transient information T(38, 39). Furthermore, EIBs are processes that persist in the environment and that other agents inherit and exploit, without duplicating the cognitive effort that built them. In this way, animals and human beings have used their environment to expand the efficiency of their bottlenecks.

#### Animals’ material environments are EIBs

Animals’ constructions are adaptations that become integral components of the organism’s phenotype (77). Natural selection acts not only on bodies but also on the external structures that organisms build (78). Many animals shift cognitive load into their surroundings by making burrows, nests, dams, and bowers. These act as EIBs that render tractable the synchronization of information relevant to survival and reproduction. For example, the burrow of the long-finned goby (*Valenciennea longipinnis*) is typically seen as regulating water flow, egg protection and oxygenation (78). Computationally, it converts an unpredictable hydrodynamic environment into a stable and predictable one. The burrow filters environmental variability–the action observation space— so that only a narrow range of relevant states reaches the fish. A spider web converts complex, high-dimensional motions of flying insects into localized vibrations. By building its web, it refines its channel capacity, increasing the action-observation types used to determine the position of an insect. Thus, the spider too transforms a difficult perceptual problem into a tractable one.

Evolution repeatedly rediscovered this neat solution. It is often easier to simplify the environment than to make organisms more complex. Furthermore, the resulting EIBs can also be inherited and exploited. From the ecological niche construction theory perspective, species actively modify their shared environment constraining the selective pressures for evolution (79, 80). A beaver dam transforms the river in a local environment, making water flow easier to predict, hence reducing hydrodynamic uncertainty. Other species inherit these EIBs, constraining their available actions and thereby creating selection pressure for inference strategies that better comply with these constraints (80, 81).

Of course, not all situations end positively. AIBs can also fail when they become too local, capturing regularities that no longer generalize. Rituals, bureaucracies and monetary systems, achieve remarkable reductions in complexity by compressing heterogeneous realities into tractable rules, categories, tokens, or measures. Overfitting occurs when an EIB becomes increasingly optimized to the compressed regularities generated within its own niche, while losing sensitivity to the wider environmental structure it originally evolved or was designed to make tractable.

Financial markets are an extremely good example of EIB overfitting because prices are simultaneously representations of expectations about the world, and causes of subsequent market behavior that changes that world. This recursive nature of the market makes that the regularities encoded by these representations cease to generalize, the EIB continues to organize behavior according to a model that no longer has a direct bearing on the world. In doing so, it systematically privileges what can be easily represented while obscuring qualities that resist quantification, such as ecological resilience, tacit knowledge, or long-term social value. As decisions become increasingly guided by the compressed representation itself, the metric, category, or token ceases to track reality and instead becomes the target of optimization, preserving compression at the expense of adaptation.

By contrast, *Hostile scaffolding* (82) names an insidious case, in which an environment (such as some forms of social media) exploits the agent’s reliance on external structures, serving the interests of the party who designed it. The user is led to believe the tool is helping them, when in truth its design works against their interests. By disguising manipulation as assistance, it constrains the user’s choices toward a goal (such as getting ‘likes’) that belongs not to them, but to whoever designed and profits from the device. The machinery appears benign: familiar, intuitive, even helpful. Yet, deep down, every apparent reduction in cognitive load is engineered to steer the user further from genuine autonomy and toward the designer’s ends (see also: (83)). The question then becomes who controls the bottlenecks controls the information that enters our internal models of the world and, ultimately, who we are.

#### Cultural EIBs

Human culture itself acts to produce EIBs, selectively amplifying certain forms of information while suppressing others. Examples can be found in architecture and artificial landscapes that channel situated perception and action; roads that channel movement; maps that compress geographic information; calendars that do the same for timing and seasonal variations, increasing structure temporal cycles and temporal depth; and language and writing which confine expression into discrete concepts and symbols. Rituals, myths, and institutions likewise shepherd social interactions (Fig. 2), while the creation of social norms itself plays a similar role, constraining behaviors and simplifying prediction and inter-agent interpretability. Other interesting examples of cultural EIBs can be found in SI.

Human way-finding (84) affords a simple example. Wayfinding is the ability to reliably reach a destination in spatially extended environments. Navigation across cultures relies heavily on the use and creation of environmental cues, systematized knowledge, visualization techniques, and cognitive artefacts. Traditional navigators do not solve navigation solely through internal computation. Instead, they exploit culturally transmitted external artifacts such as stick charts, songlines, celestial schemas, tactile maps, and other culturally transmitted resources that offload part of the inferential burden of constructing and maintaining spatial representations of the world. From the perspective of the TC framework, these technologies are EIBs that solve the CII related to the problem of wayfinding. As we showed in our example (Fig. 4), for correct spatial navigation agents need to build tractable egocentric and allocentric representations. Thus, rather than requiring all navigational computations to be performed internally, cultural artefacts, landmarks, monuments, and other EIBs transform a high-dimensional inference problem into a tractable one (85, 86, 25).

Importantly, cultural differences do not simply reflect how effectively AIBs resolve the CII, but also which aspects of the world different cultures engage with. In fact, different AIBs may select different subsets of information, while multiple AIBs can operate on the same subset (Fig. 2). Thus, cultural variation can reflect both different solutions to similar problems and/or adaptation to different epistemic niches.

More generally, wayfinding can be interpreted as a foundational form of cognition. Before humans navigated abstract conceptual spaces, they first had to navigate physical ones (87, 88). The same logic may later have extended to concepts and language themselves. Just as navigational artefacts made physical environments tractable, conceptual and linguistic systems function as internal AIBs that make increasingly complex domains tractable (Fig. 2). In this view, EIBs may have freed inferential resources for the development of higher-order conceptual, linguistic, and symbolic representations

## Conclusion

Adaptive information bottlenecks characterize a wide continuum of neuro-ecological artifacts spanning nervous systems, animal body-plans and physiology, and socio-cultural products. This reveals a hidden continuity that speaks to the very heart of what cognition is and how it functions. The picture on offer can also be seen as a kind of spatio-temporally generalized version of the ‘extended mind’ hypothesis (89) in which externalized bottlenecks are environmental transformations that function analogously to information bottlenecks in the brain, but are realized in the environment to solve CII.

There is a direct fit also with work in active inference that formalizes intuitions from early cybernetics. Cybernetics established that regulation depends on information (19) (Ashby’s Law of Requisite Variety), while active inference formalizes homeostasis and allostasis as an inferential process in which organisms maintain viability by minimizing uncertainty through action (9), guided by informational boundaries such as Markov blankets (90). For example, the cell membrane functions as a physical, chemical, and informational boundary that regulates exchanges to preserve homeostasis (90). Within this framework, allostasis— “stability through change” (91) —is not limited to internal regulation but extends to organism–environment interactions to reduce uncertainty (9). In this sense, regulation under active inference is distributed across the brain, body, and world through internal and external AIBs that simplify the space of possible actions.

From a holistic perspective, life progressively externalizes information processing. At first, cells evolve membranes, then animals evolve webs, nests, burrows, mounds, bowers, dams, etc. Thanks to cumulative material culture, humans then evolve houses, villages, monuments, roads, maps, writing, cities, archives, digital technologies, and generative AI. Each innovation reduces inferential burden, exploits new hidden regularities and enhances predictability, by the agent or the collective, to maintain viable states of life. Therefore, Tractable Compression can be analyzed at different levels of granularity where AIBs may characterize the behavior of individual organisms, coordinated groups, or entire cultures. Each level gives rise to its own repertoire of solutions to the CII. Thus, different species, human cultures, and animal social systems can all be viewed as alternative individual or collective strategies for addressing the CII.

Human, animal and cultural history, all reflect the need to solve the fundamental challenge of information intractability by the progressive construction of AIBs that extend allostatic control further and further into the world.

## Materials and Methods

### Computational mechanics model

We modeled two source process as a discrete-time, three-state Markov system with hidden states A, B, and C. The hidden-state transition matrix for both processes were:

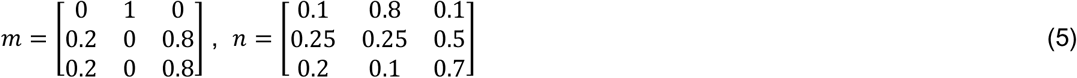

All simulations were initialized in state A and generated for t=10^6^ steps using a fixed random seed. Holding the latent dynamics fixed across conditions to isolate how alternative observation channels constrain the block entropy profile.

The true entropy rate of the underlying hidden Markov chain h_μ_ was computed from the stationary distribution *π_i_* and then evaluating

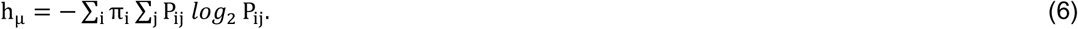

where *π_i_* is the stationary distribution (obtained by solving 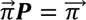, and *P_ij_* is the transition probability from state *i* to state *j*. This equals the conditional entropy *H*(*X*_*t*+1_ ∣ *X_t_*) under stationarity ((18), Theorem 4.2.4).

### Channel capacity

Because some hidden transitions had zero probability under the chosen transition matrix, the effective channel capacity depended on the number of emitted symbols actually observed in the simulated sequence (*A_obs_*). Capacity was therefore computed empirically as,

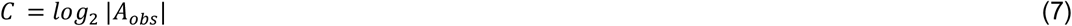

where |*A_obs_*| is the number of unique observed symbols in a given condition.

### Block entropy and finite-length entropy rate

For each emitted symbol sequence, block entropy was computed for block lengths *L* ≥ 1 where *H*(0) = 0. We applied a sliding window of length L that advanced one measurement at a time from left to right. Each window produced a contiguous block of length L, and the frequency of these blocks was used to computed block entropy as

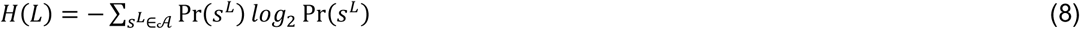

The apparent entropy rate 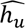 was computed as

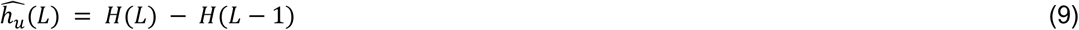

where ℎ(0) = *C* following (39). The final 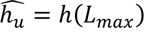, is understood as the local synchronization achieved for each different observation channel scenario. This allowed to compare the local finite estimated entropy rate 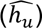 for each observation channel against the true entropy rate of the source (ℎ*_u_*).

The predictability gain was computed as

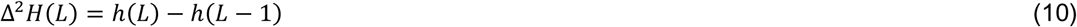

for each condition, the terminal finite-length estimate was used as the asymptotic estimate of the estimated entropy rate. Based on this estimate, we computed finite-length excess entropy

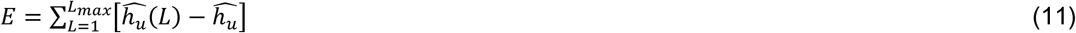

where, E (bits) measures how much information the past carries about the future. This leads to the notion of the cost of amnesia: if an agent loses access to its past, how much additional randomness will the environment exhibit.

We also computed a finite-length estimate of the transient information:

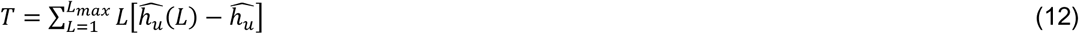

where T denotes bits x symbols and gives an effective measure of synchronization cost by penalizing the need for long observation blocks required for synchronization.

### The information bottleneck method

The Information Bottleneck (IB) was computed using the Blahut-Arimoto algorithm implementation (from https://www.mathworks.com/matlabcentral/fileexchange/65937-information-bottleneck-iterative-algorithm by Shabab Bazrafkan) for each channel and process. For each channel, the IB was resolved for a codeword of size |T| = 10. Variations in different t values did not significantly affect the final solution, so a high value was selected to efficiently summarize the results. Convergence was determined using a Jensen–Shannon divergence threshold of ε = 10⁻¹⁰.

All code was executed in Python and MATLAB and can be downloaded at https://github.com/Rockyneuron/tractable_compression

## Acknowledgments

This project was supported by a European Research Council, Synergy Grant (XSCAPE) ERC-2020-SyG 951631. We would like to thank all our colleagues and the technical staff at INCIPIT-CSIC for their support and assistance throughout this work.

## Author Contributions

Conceptualization: A.V.P., L.M.M.; Methodology and software: A.V.P.; Formal Analysis: A.V.P., A. Co., A.C.G.; Visualization: A.V.P., A.Co.; Supervision: L.M.M., F.C, A.Cl.; Writing—original draft: A.V.P. L.M.M; Writing—review & editing: All authors

