## Supplementary Information for "Compression through internal and external information bottlenecks resolves information intractability across brains, bodies and worlds"

### Supporting Information Text

### Information Bottleneck Compression-Relevance tradeoff

The IB method is a variational approach to find a compressed representation of X, which is Z, that selects relevant information to predict another signal Y(1). So, the problem comes to find a $\mathrm{p(z|x)}$ that minimizes the functional

$$\mathrm{F[p(z|x)] = I(Z;X) - \beta I(Z;Y)} \quad (1)$$

Where $\mathrm{I(Z;X)}$ measures the amount of information about X retained by Z, whereas $\mathrm{I(Z;Y)}$ measures the relevance of Z for predicting Y, and β > 0. The parameter β controls the trade-off between compression and prediction. Minimizing $\mathrm{I(Z;X)}$ relates to optimally compressing X for the prediction of Y. Maximizing $\mathrm{\beta I((Z;Y)}$, relates to how much predictive power Z will have. The predictive information preserved by Z is bounded by the information that X itself carries about Y, where I(Z;Y) ≤I (X;Y) (2). Thus, the IB method characterizes a set of relevance-compression solutions, each retaining different amounts of information about X while preserving different amounts of relevance for Y.

### IB trade-off between the past and future information of the channel

To quantify how much of the observation stream is predictive of what comes next, we applied the Blahut-Arimoto algorithm (see methods) to the empirical joint distribution of past and future observation windows. Let $x_{1:n}$ denote the observation sequence for a given channel. For each time index $t$, we defined a past window of length $L$ and a future window of length $K$ as

$$X_t^{(L)} = (x_{t-L}, \dots, x_{t-1}), \qquad Y_t^{(K)} = (x_t, \dots, x_{t+K-1}) \quad (2)$$

We used $L = 2$ and $K = 1$, so that $X_t^{(2)} = (x_{t-2}, x_{t-1})$ and $Y_t^{(1)} = x_t$

Thus, $X$ summarizes the immediate past and $Y$ the immediate future, including the present symbol when $K \geq 1$. We then estimated the joint distribution $P(X, Y)$ from the observation stream by counting co-occurrences and applied the Blahut-Arimoto algorithm to compress $X$ (the past) while preserving relevant information about $Y$ (the future).

### The CII of wayfinding or navigating the environment

In the main text, we use spatial navigation and GPS systems to illustrate the relocation of navigational burdens into externalized distributed infrastructures. Here we expand this argumentation to provide cultural and historic examples of how the EIBs mechanisms solve the CII problem by tractable compression. Next, we extend the reference to wayfinding and spatial navigation and discuss how the same problems are solved, providing examples from prehistoric and historic monumentality.

Visual navigation requires the construction of coherent egocentric and allocentric representations from information distributed across the environment, body, and nervous system. In this section, we examine how this process can be understood as a sequence of coupled AIBs operating across multiple scales. As an agent becomes entangled with the available information, its initial position in the environment establishes the first information bottleneck. That bottleneck determines the information accessible from the agent's location and therefore already reduces the dimensionality required to resolve the task. Thus, the agent's position in the world can be regarded as the initial Adaptive Information Bottleneck (AIB) that already constrains the action-observation space (AOS).

Additionally, the agent can get more precise information of its location using a GPS. The flow of information from a satellite-radiosignal-mobile device (process-emitter-receiver) is an environment only AIB (i.e., environment-environment-environment, meaning that all three components of the IB, brain-body-environment, are located in the environment). Each satellite emits its position to earth allowing a mobile device to make out its precise location in the world.

Once our mobile device has determined our location, the next stage involves an information bottleneck that straddles the environment, the body, and the brain. Visual information is reflected from the environment and enters the eye, projecting onto the retina through the cornea, lens, and ocular media. The size, geometry, and optics of the eye limit the amount of environmental information sampled at any given time, thus establishing the spatial and temporal boundaries of the retinal image.

The next stage in this continuous flow of information includes a new bottleneck located entirely within the nervous system. The human retina contains approximately 100 to 120 million photoreceptors, but only 1 to 1.5 million ganglion cells, whose axons constitute the optic nerve (3, 4). Consequently, the information captured by the photoreceptor layer is anatomically compressed by almost two orders of magnitude before reaching the brain. From an information theory perspective, this compression is especially significant, indicating that selective sampling and inference is not merely a cortical phenomenon but a fundamental information processing strategy already present at the earliest stages of the visual pathway, in particular, and sensory transduction, in general (5, 6). This indicates that every sensorial perception (not only visual) is not primarily a process of representing the world, but of progressively selecting, compressing, and stabilizing behaviorally relevant information (7, 8).

Visual navigation, therefore, requires both the acquisition of environmental information and the making of manageable inferences. Satellite navigation systems, ocular optics, retinal circuits, and the cognitive processes involved, act as mechanisms that selectively restrict the flow of information, reducing an otherwise overwhelming sensory flow to a manageable set of states for action. Monuments, maps, roads, landscapes and other cultural artifacts and objects (including portable ones) can function as higher-order bottlenecks, in this case simplifying inference at the level of behavior, navigation, and social coordination, rather than at the neuronal level.

### Extending examples of external information bottlenecks

**Monumentality and European Atlantic megalithism**. Prehistoric megalithism provides a particularly illustrative example of EIB. Emerging in the Neolithic and persisting for over two millennia (ca. 6700–4200 BP) megaliths represent are among the earliest forms of monumental architecture. Although earlier examples of monumentality, such as Göbekli Tepe in Turkey (ca. 11600-10200 BP) (9), are known, megalithism marks one of the first traditions of monument construction, even before the pyramids (10). Thousands of megalithic monuments were built across Atlantic Europe, from North Africa to southern Scandinavia (11). Typically, these structures consist of large stone-built funerary chambers (the term megalith literally means “big stone”) enclosed within earthen and stone mounds. These structures range from measuring just few meters in diameter, to over 90 meters, such as Newgrange in Ireland, a UNESCO World Heritage Site (12). Monuments, and particularly megaliths, are characterized by four defining properties, relevant for understanding EIBs: as large-scale artificial constructions, they are *spatially visible*, *temporally enduring* and require substantial *planning and coordinated labor.* Their significance, however, lies in the fact that megaliths actively structure human spatiotemporal experience and engagement with the environment (13, 14). In terms analogous to the retina, from an EIB perspective, megaliths can be understood as shaping an information space: a structured subset of environmental information with which agents attempt to synchronize.

This process is particularly significant because it suggests that the domestication of space and time preceded the actual domestication of the broader environment. Illustrating how humans did not merely adapt but actively transformed their environment to cognitive resources, shaping their minds while making monuments. Megaliths successively guide visual attention, structure movement through space, and ultimately create cognitive maps that impose cultural structure on the world (15).

By channeling attention toward specific locations, horizons, routes, and seasonal events, monuments build landscape, organize social temporality and shape perception, memory, and interaction. In this sense, they introduce environmental regularities that modulate channel capacity and reduce the range of relevant information, constraining the actions required to navigate social and physical environments.

Over time, their topological disposition of megalithic monuments determines which hidden environmental states are revealed and which places become salient through repeated engagement. This constitutes a form of counterfactual sampling: instead of exploring the landscape uniformly, agents selectively sample the most informative ones.

Across generations, these material structures become stable cognitive and social resources. By scaffolding memory, orienting collective attention, synchronizing rituals, and stabilizing expectations about the world, they function as durable externalized predictive infrastructures. In this way, they reduce the inferential burden of coordinating action across space and time, functioning at the level of individuals, social groups or even strangers in the same landscape. Thus, by guiding attention, the number of actions required for egocentric and allocentric positions in the environment decreases (selective sampling).

Rather than requiring each generation to reconstruct territorial, cosmological, or seasonal knowledge from scratch, monuments embed relevant information in accessible material form. Thus, they do not merely reflect social organization, but they actively contribute to it by reshaping informational and cognitive ecologies within which people act. By stabilizing long-term regularities and making future events more predictable, they increase temporal depth, enabling planning, delayed reward, seasonal coordination, and transgenerational memory. As persistent materialised processes, megaliths constitute inherited EIBs that subsequent generations can exploit without having to invest the original labour and cognitive effort. They thereby function as systems of transgenerational cognitive offloading, demonstrating how human beings, as cultural animals, progressively use their environment to expand the efficiency of their bottlenecks and to externalise more and more cognitive load into their surroundings.

The implications of this extend beyond Atlantic megalithism. Monumental constructions across history share similar properties. From Atlantic megalithism to modern urban settings (from the royal palaces and gardens of Versailles in France -XVII century- and Alcalá de Henares in Spain -XVI to XVIII century, to the modern urban planning of Washington, DC, and Paris) as well as prehistoric pyramids or contemporary skylines, embodying power and prestige, monumentality has operated in a consistent manner.

**Catholic and Lutheran churches.** Another example of EIB can be found in the distinct liturgical apparatus of Catholic and Lutheran churches. Although these two religious spaces serve a similar purpose, namely synchronizing individuals with God and the community, they achieve it through different means.

In terms of channel capacity, Catholic spaces reduce action complexity through imagery (saints, icons) that refer to particular passages of the Bible, whereas Lutheran settings prioritize direct textual guidance. From the perspective of counterfactual sampling, while both maintain a similar spatial organization, they differ in the nature of the material tools they employ. This, in turn, changes the symbolic focus. Albeit through different approaches, ultimately both traditions reduce the number of actions required for synchronization.

**Frankenstein and RAG-Large Language Models.** Mary Shelley's Frankenstein (1818) is meant to be a story about the dangers of scientific hubris, but it also contains one of literature's most sophisticated depictions of cognitive development and how minds are shaped by their environments full of externalized information bottlenecks. The Creature's gradual acquisition of language and literacy unfolds through a handful of social encounters, books, and written documents, offering an early literary account of how EIBs can structure the final inference strategies an agent can develop.

In the novel, the creature first learns to speak, and language in general through observation, imitation, and social inference. After fleeing from Victor Frankenstein, the Creature eventually hides in a hut attached to the cottage of the De Lacey family, an elderly blind father and his children, Felix and Agatha. By secretly observing the De Lacey family he discovers statistical regularities between events and vocalizations, some structure in the otherwise apparently random process of vocal communication that makes him realize that certain sounds correspond to people, others to objects, and yet others to actions.

At some point, watching the De Laceys read and write, he realizes that writing is a symbolic system that somehow represents and preserves speech. The novel implies that he gradually learns letter-symbol correspondences and narrative, effectively self-teaching himself to read. The following step in his learning process came when he found four books: Ruins of Empires by Comte de Volney, which provides the creature with his first basic instruction on human history, social structure, governments, and religion; Paradise Lost by John Milton, perhaps the most important, because it serves to shape his moral and existential framework, and uses it to understand his relationship with his creator, deeply relating himself to the outcast Adam; Plutarch's Lives of Illustrious Greeks and Romans, from which he learns about human politics and societal virtues; and, finally, The Sorrows of Young Werther by Goethe, which helps him understand complex human emotions and passions. The final step in his learning process was finding Victor Frankenstein's journals which contained an account of how he was created, how Victor regarded him, and why he finally abandoned him. Reading these journals radically changes the Creature's self-model and his moral compass.

In a recent article, Martin Puchner, looked at Frankenstein in terms of Large Language Models (LLM) and the framework of Retrieval-Augmented Generation (RAG) that connects LLMs to external knowledge bases (16). Instead of relying solely on previously learned training data, RAG pull relevant, up-to-date information from selected external sources first, allowing the LLMs to generate highly accurate, fact-based responses, injecting context, and drastically reducing the rate of hallucinations. Puchner suggested that this is very similar to the Creature's learning process, first going through an LLM phase in which he learns to speak and to use language as a tool observing the De Laceys; then going through a RAG phase, when he reads just four books that will shape his subsequent behavior.

The learning trajectory of Frankenstein's Creature can be interpreted through the framework of TC by examining changes in channel capacity, counterfactual sampling, and selective sampling. Upon discovering the De Lacey household, the Creature enters a structured informational environment that substantially reduces channel capacity: the complexity of learning human social behavior is constrained by a stable setting where language, literacy, and social interaction are continuously enacted. This environment enables efficient counterfactual sampling, as the Creature explores and compares multiple observational and behavioral possibilities (communication, reading, and social interaction) within a bounded context. Over time, the acquisition of language and higher-order concepts supports increasingly efficient selective sampling, allowing the Creature to navigate the world with reduced uncertainty and enhanced predictive capacity. Together, these processes reduce transient information, i.e., the informational cost required for synchronization with social reality.

What is remarkable is that The Creature's entire cognitive development depends only on a tiny set of external information channels that constrain and shape the very space of possible thoughts available to him. He does not merely learn from the books, his expectations, moral concepts, and counterfactual imagination become structured by the very small collection of cultural sources through which selected information about an unfathomable world reaches him. This is essentially our current definition of an efficient externalized bottleneck; the four books allowing the Creature to access new hidden states from which to build an internal model of the world that is vastly larger than his direct experience would allow.

**Social and cultural AIBs.** Social and cultural bottlenecks all likely operate this way in their specific historic contexts. Social institutions and norms, administrative regulations and hierarchies, educational programs, and political and legal systems, constrain how agents behave on a given place and time; they stabilize particular attentional habits, structure the range of expectations, define inferential strategies, and, as in the case of Frankenstein's creature, limit the space of possible thoughts available.

Cultural, scientific, and creative activities operate in the opposite direction, contributing to social and cultural evolution by increasing counterfactual bandwidth and capacity, effectively expanding the inferential landscape of future generations. Yet there is a darker side to this process when mass-produced entertainment, from music and cinema to the internet and social media algorithms, becomes so heavily curated that it primarily serves to standardize and westernize human experience, promoting consumerism, fueling misinformation, and increasing ideological extremism and affective polarization (17, 18).

**Cognitive enhancement and neuroprosthetic devices.** According to the tractable compression framework (TC), cognition is not fundamentally limited by the number of neurons or synapses, but by the coupled AIBs operating across body, brain and world. Neuroprosthetics and cognitive augmentation technologies can therefore be understood as attempts to modify, bypass, or create new forms of AIBs.

The history of cognitive enhancement is thus mostly the history of external information bottlenecks (EIBs) engineering. The greatest cognitive revolutions often come not from changing brains but from altering EIBs. Writing, for example, did not increase the brain's volume, it externalized memory; maps did not increase working memory, they externalized spatial inference; and mathematics did not arguably increased IQ, it simply externalized complex computations.

Future neurotechnology follows the same logic. A retinal implant is not just a sensory replacement, it is an engineered information bottleneck where millions of photons are compressed into a limited number of stimulation channels. Therefore, this is the same design problem that evolution solved in the retina, what information should be discarded, what should pass through, and at what temporal scale? From this perspective, neuroprosthesis are, indeed, the artificial equivalent to anatomical information bottlenecks (19).

Today, our internal models of the world are already partially externalized through the use of notebooks, smartphones, search engines and AI agents, social networks, GPS, and even scientific papers such as this one. One can imagine the future of neural interfaces making these connections far tighter resulting in internal models that become hybrid biotechnological systems. Looking back to Mary Shelley's Frankenstein, a cautionary note is in order; the Creature creates his model of the world from a handful of externalized bottlenecks, future augmented humans may do something similar through neural interfaces connected to large external knowledge systems that are continuously accessible. The question then becomes who controls the bottlenecks controls the information that enters our internal models of the world and, ultimately, who we are.

### Figures

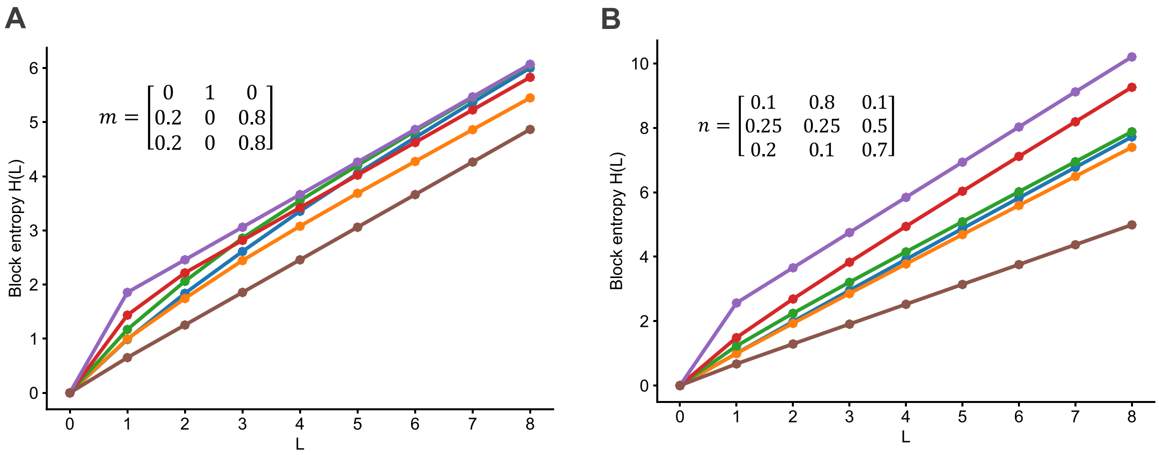


**Fig. S1**. **Block entropy profiles of the AIB model**. **A**. Block entropy of the m process. **B**. Block entropy of the n process.

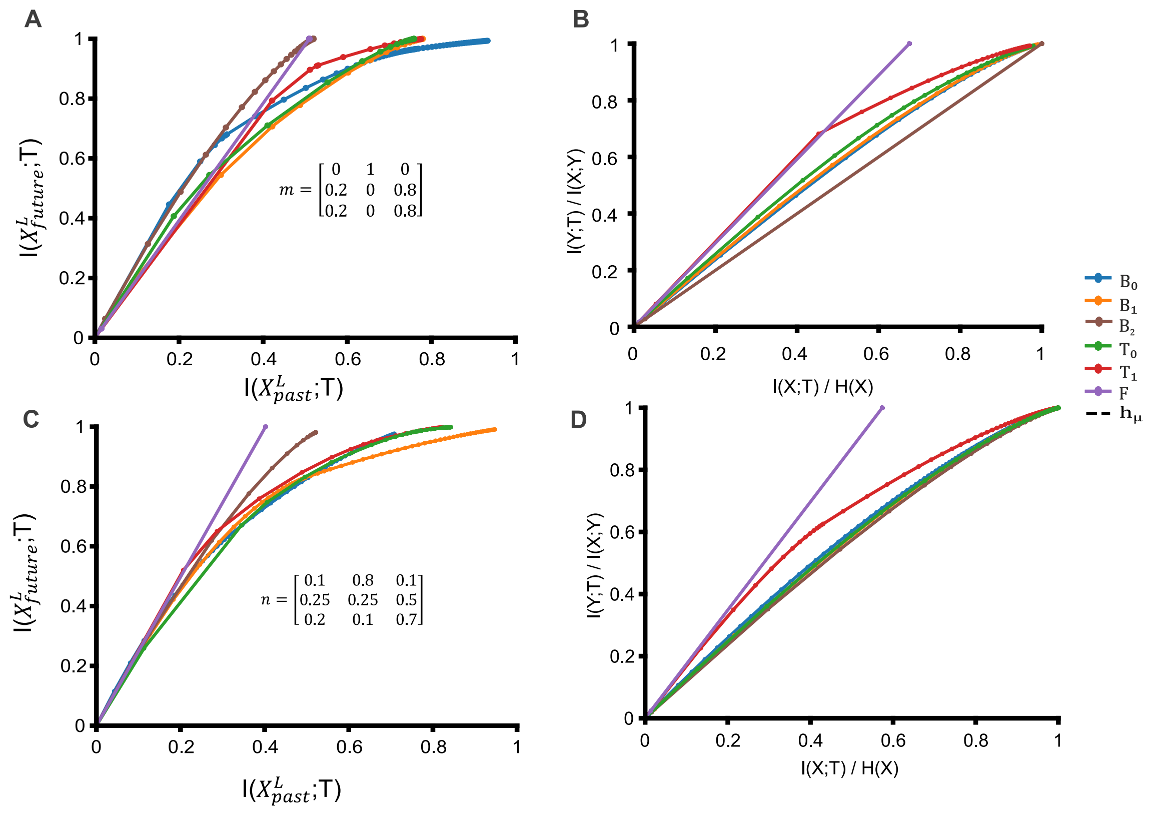


**Fig S2**. **Normalized Information Bottleneck trade-off profiles of the AIB model**. **A, C**. Normalized IB profiles relating compression to predictive future and past information across AIB channels for the m and n processes, respectively. **B, D**. Normalized IB profiles relating the observation channel (X) and hidden state process (Y) for m and n processes, respectively. Normalization allows comparison of channel efficiency independently of absolute information content.

### Tables

| | $E$ | $T$ | $\widehat{h_u}$ | $h_\mu$ | error | rel error | $C$ |
|---|---|---|---|---|---|---|---|
| $B_0$ | 0.9154 | 2.1984 | 0.6356 | 0.6016 | -0.0340 | -0.0565 | 1 |
| $B_1$ | 0.7543 | 1.3785 | 0.5867 | 0.6016 | 0.0149 | 0.0248 | 1 |
| $B_2$ | 0.0488 | 0.0492 | 0.6021 | 0.6016 | -0.0005 | -0.0009 | 1 |
| $T_0$ | 1.1843 | 2.4136 | 0.6095 | 0.6016 | -0.0079 | -0.0131 | 1.5850 |
| $T_1$ | 1.0115 | 1.1890 | 0.6021 | 0.6016 | -0.0005 | -0.0008 | 1.5850 |
| F | 1.2532 | 1.2539 | 0.6021 | 0.6016 | -0.0005 | -0.0008 | 2.3219 |

**Table S1**. **Block entropy measures for different AIB under the first hidden-state transition matrix**. For each observation channel, the table reports the excess entropy $E$, transient information $T$, estimated entropy rate $\widehat{h_u}$ true entropy rate $h\mu$, error, relative error, channel capacity $C$. Signed error was calculated as $h_\mu - \widehat{h_u}$ , such that positive values indicate underestimation and negative values indicate overestimation. Relative error was calculated as $(h\mu - \widehat{h_u})/h\mu$.

.

| | $E$ | $T$ | $\widehat{h_u}$ | $h_\mu$ | error | rel error | $C$ |
|---|---|---|---|---|---|---|---|
| $B_0$ | 0.1108 | 0.2557 | 0.9581 | 1.2065 | 0.2483 | 0.2058 | 1 |
| $B_1$ | 0.1262 | 0.2237 | 0.9170 | 1.2065 | 0.2895 | 0.2399 | 1 |
| $B_2$ | 0.0428 | 0.0437 | 0.5828 | 1.2065 | 0.6237 | 0.5170 | 1 |
| $T_0$ | 0.3740 | 0.5383 | 91.0080 | 1.2065 | 0.1984 | 0.1645 | 1.5850 |
| $T_1$ | 0.5055 | 0.8914 | 1.1548 | 1.2065 | 0.0517 | 0.0428 | 1.5850 |
| F | 1.5161 | 1.6739 | 1.1949 | 1.2065 | 0.0115 | 0.0096 | 3.1699 |

**Table S2**. **Block entropy measures for different AIB under the second hidden-state transition matrix**. For each observation channel, the table reports the excess entropy $E$, transient information $T$, estimated entropy rate $\widehat{h_u}$ true entropy rate $h\mu$, error, relative error, channel capacity $C$. Signed error was calculated as $h_\mu - \widehat{h_u}$ , such that positive values indicate underestimation and negative values indicate overestimation. Relative error was calculated as $(h\mu - \widehat{h_u})/h\mu$.
